# RNase R Controls Membrane Fatty Acid Composition in *Streptococcus pneumoniae*

**DOI:** 10.1101/2023.03.21.533657

**Authors:** André Filipe Alípio, Cátia Bárria, Vânia Pobre, Rita Matos, Mónica Amblar, Cecília Maria Arraiano, Susana Domingues

## Abstract

Previous studies on RNase R have highlighted significant effects of this ribonuclease in several processes of *Streptococcus pneumoniae* biology. In this work we have studied the global impact of RNase R by comparing the transcriptional landscape of a deleted RNase R mutant to that of the wild-type strain, and this led us investigate specific targets affected by RNase R. RNA-Seq showed that RNase R deletion affects transcripts from several different biological processes. Of particular interest, elimination of RNase R results in overexpression of most of the genes encoding the components of type II fatty acid biosynthesis (FAS-II) cluster. We demonstrate that RNase R governs the turnover of most of genes from this pathway, affecting the outcome of the whole FAS-II cluster, and leading to an unbalanced membrane fatty acid composition. Our results show that the membrane of the deleted strain contains a higher proportion of unsaturated and long-chained fatty acids than the wild type strain. This leads to a higher fluidity of the Δ*rnr* mutant membrane, which is probably related with the increased sensitivity to detergent observed in this strain. We demonstrate that RNase R expression is induced in cells challenged with H_2_O_2_, which is suggestive of a role for this ribonuclease on the regulation of membrane homeostasis under oxidative stress. Reprogramming of membrane fluidity is an adaptative cell response crucial for bacterial survival in constantly changing environmental conditions. The fact that RNase R controls the expression of several essential genes to the fatty acid synthesis unveils a new important function of this enzyme.

## Introduction

*Streptococcus pneumoniae* is an important human respiratory pathogen that causes a variety of serious diseases. It is an opportunistic pathogen commonly found in the nasopharynx flora of healthy individuals. Pneumococcal colonization usually occurs on the mucosal surface of the nasopharynx during childhood and in most cases persists asymptomatically into adulthood [1]. Transition from asymptomatic colonization to disease occurs when *S. pneumoniae* migrates to other tissues of the human body. This bacterium often colonizes the lungs being the leading cause of bacterial pneumonia. Colonization of other body tissues may give rise to meningitis, otitis media and sepsis. The pneumococcus affects especially young children, the elderly and immunocompromised individuals. Diseases caused by pneumococci constitute a major public health problem. In 2017, the World Health Organization included *S. pneumoniae* in the list of 12 priority pathogens that urgently require new antimicrobials. The emergence of antibiotic-resistant strains has conferred this bacterium the status of superbug.

Normal disease progression to the lung or bloodstream exposes *S. pneumoniae* to various stress conditions and fast adaptation to these challenging situations is a crucial step to bacterial survival. Adjusting the RNA amount at the post-transcriptional level, ensures a faster and less energy-demanding cell response [2]. Ribonucleases (RNases), by controlling the amount of every transcript in the cell, are key effectors in bacterial adaptation and survival under stress [3]. Despite their important roles, knowledge on pneumococcal RNases is still in its infancy, with only a small number of recent studies. RNase R, PNPase and RNase Y were all shown to affect *S. pneumoniae* virulence [4, 5]. PNPase seems to act on virulence through its interaction with noncoding RNAs [5, 6]. In a recent study RNase YhaM was identified as a stabilizer of noncoding RNAs involved in pneumococcal competence [6]. The present work is focused on RNase R, an exoribonuclease that degrades RNA in the 3’ to 5’ direction in a processive and sequence-independent manner, being the only 3’-5’ exoribonuclease able to degrade RNAs with extensive secondary structures [7-9]. RNase R belongs to the RNase II/RNB family of enzymes, which is present in all domains of life [7]. In *S. pneumoniae* RNase R is the only known member of the RNB family, suggesting a major role in this microorganism. Our work on RNase R has already elucidated very interesting roles of this enzyme on the pneumococcal biology. RNase R responds to cold-shock and regulates the expression of the *trans*-translation effector SmpB, which in turn affects RNase R levels [10]. We have also established RNase R as a ribosome dissociation factor, which affects the amount of active translating ribosomes by controlling the level of ribosome dissociation factors. This has a global impact on protein synthesis and affects cell viability [11]. In addition, our work on the effect of RNase R on *S. pneumoniae* virulence has shown that RNase R interferes with the host immune response, probably by affecting the ability of the pneumococcus to internalize *Galleria mellonella* hemocytes [4].

The overall body of evidence suggests a global role of RNase R on the physiology of this bacterium with repercussions on the *S. pneumoniae* ability to cause infection. Therefore, in this work we have performed a global transcriptomic analysis with RNA sequencing to determine the most significant differences between the wild-type (WT) strain and an isogenic mutant deleted of RNase R (Δ*rnr*). To our knowledge this is the first RNA-seq study focusing on the role of RNase R in *S. pneumoniae*. The data presented here shows an important role of RNase R in the control of the fatty acid composition and therefore membrane fluidity. We demonstrate that RNase R governs the turnover of genes from the Fatty-Acid (FA) biosynthesis pathway, leading to an unbalanced level of membrane FA. The possible implications of the altered membrane FA composition on resistance to detergent and oxidative stress are further explored.

## Results

### Screening of RNase R-regulated genes via RNA-Seq

Transcriptomic analysis was performed to better understand how RNase R affects gene expression in *S. pneumoniae* clinical isolate TIGR4. We compared the transcriptome between Δ*rnr* mutant previously constructed [10, 12] and the WT by performing RNA-Seq of total RNA. We found 257 transcripts that had high expression values in either strain and more than 0.5 of log_2_FC between the two strains (Table S3 in Supplementary Material). These transcripts were charted in a heatmap to show their normalized expression values and comparison between the WT and the RNase R deletion mutant (Figure 1A). Most of these transcripts belonged to the ABC transporter functional category. The second most representative category was the PTS system. Interestingly, we also found two oxidative stress genes (*Tpx* and *dnaK*) that were upregulated in the Δ*rnr* mutant when compared to the WT.

**Figure 1.**
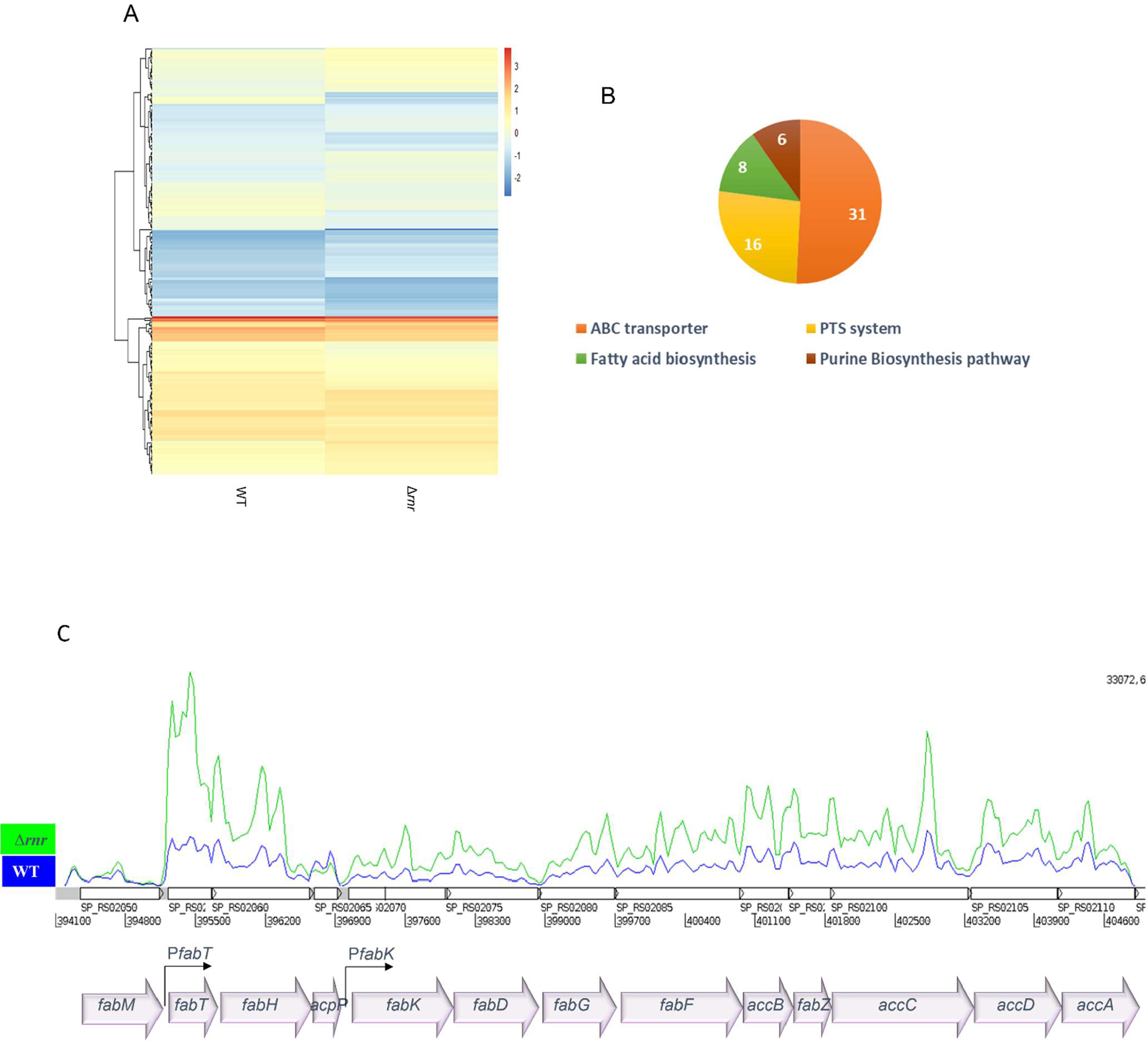
Transcriptomic analysis of RNase R deletion mutant. **A)** Heatmap of the transcriptional profile of the 257 transcripts. Hierarchical clustering was done to group genes with similar expression pattern in terms of log_2_RPKM. **B)** Representation of the functional categorization for most of the transcripts with differences between WT and the Δ*rnr* mutant. **C)** Read coverage plot of the FAS-II gene cluster. Blue line corresponds to the WT, while green line corresponds to Δ*rnr*. The y axis represents the coverage of reads, the x axis represents the gene position. The corresponding *fab* genes are represented below. Promoters P*fabT* and P*fabK* control the expression of all the cluster apart from *fabM*.

Besides these genes we also found that the purine biosynthesis pathway is also being affected by the deletion of RNase R (Figure 1B). Surprisingly, we found an enrichment of the fatty acid biosynthesis pathway. In fact, there is a striking difference in the expression of genes from the fab cluster between these strains (Figure 1C). All the genes required for type II fatty acid biosynthesis, known as the FAS-II pathway, are located in a single cluster in the *S. pneumoniae* genome, the fab gene cluster, which is represented in Figure 1C. Our RNA-Seq results show an increased expression of almost all the genes of the FAS-II cluster in the absence of RNase R, except *fabM* and *acpP*.

### RNase R controls the levels of the FAS-II gene transcripts

The influence of RNase R on the expression levels of the FAS-II gene cluster was further evaluated by Northern blot. We compared the steady-state levels of *fabT, fabH, fabK, fabG* and *fabF* by in the wild type, the Δ*rnr* and the complementing Δ*rnr*+R strain., which contains an ectopic copy of the *rnr* gene. These genes were chosen because they showed an increased expression in the RNase R deleted strain and changes in their expression levels was previously shown to result in an unbalanced level of membrane FA [13-15]. Two operons have been described within the *fab* cluster, *fab*T-*acp*P and *fabK*-*acc*A, both of which are under the control of the transcriptional regulator FabT [13, 14, 16-18]. Consistent with this, the same transcriptional unit was detected when using independently *fabT*- or *fabH*-specific probes and this unit was accumulated in the strain lacking RNase R, but not in the Δ*rnr*+R (Figure 2A). The *fabT* and *fabH* transcripts were mostly together in a co-transcribed messenger and could hardly be detected alone. The transcriptional unit containing *acpP* was also hardly visible in our conditions, which is in agreement with the idea that *acpP* either has its own promoter or its mRNA is independently stabilized (Lu and Rock 2005). The fact that *acpP* levels were not altered in the RNA-Seq results further corroborates this hypothesis. We have also analysed the expression level of *fabM* and confirmed that its level does not seem to be affected by the presence of RNase R, as expected from the RNA-Seq results (Figure S1 on supplementary material).

**Figure 2.**
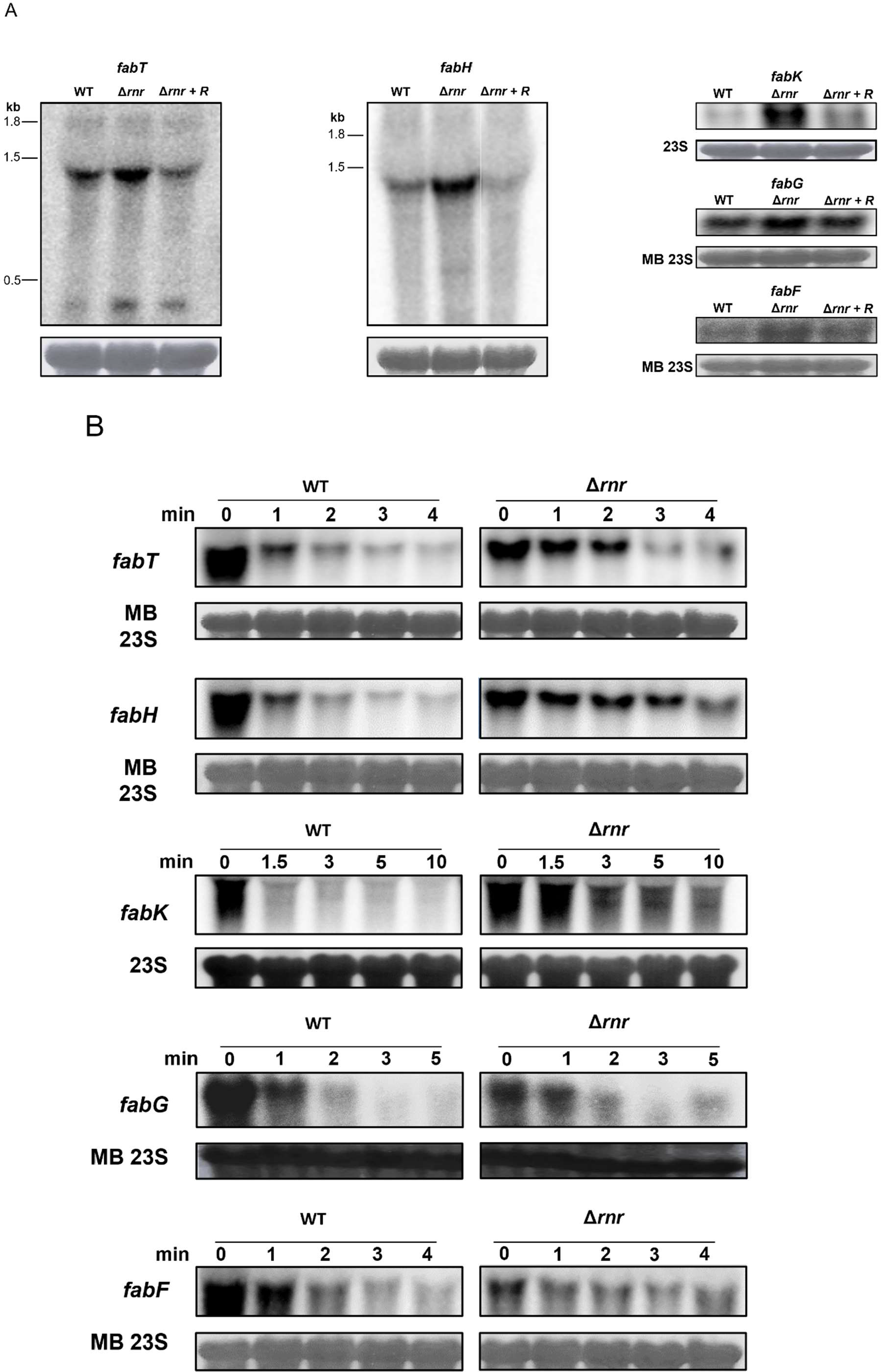
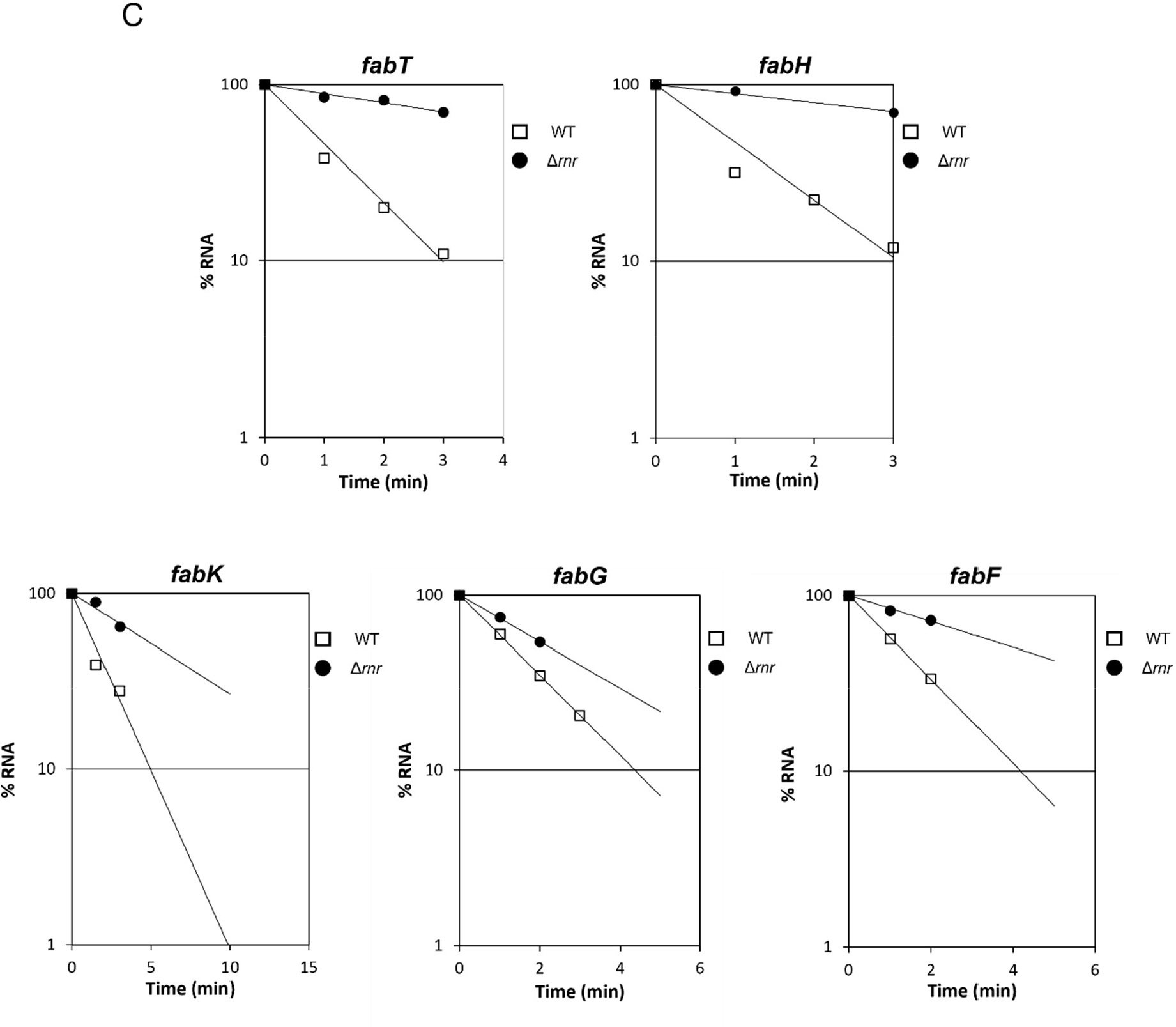
Loss of RNase R leads to accumulation of the FABII cluster genes. **A)** Northern blot analysis of total RNA samples extracted from the WT, RNase R mutant (Δ*rnr*) and Δ*rnr* ectopically expressing RNase R (Δ*rnr*+R). 20 µg of total RNA were separated on 1.5 % agarose gels. Separated RNA molecules were transferred to Hybond-N+ membranes and hybridised with specific probes. Loading controls were performed on the same membranes after stripping and probing for 23S rRNA, or by staining the membranes with methylene blue (MB) before hybridization with the probes. **B)** Northern blot analysis of the stability of the same transcripts in the WT and Δ*rnr*. Transcription was blocked by addition of rifampicin (0 min) and aliquots harvested at the indicated time-points for RNA extraction. 20 µg of total RNA were separated and blotted to Hybond-N+ membranes as described above. Gels were then hybridized with specific riboprobes. Loading controls were performed by staining the membranes with methylene blue (MB) before hybridization. **C)** Quantification of the corresponding mRNAs was done by scanning densitometry and values of the amount of RNA at zero minutes were considered 100 %. The percentage of transcript remaining after rifampicin addition was plotted as a function of time. Decay rates were evaluated by linear regression analysis. These results are the mean of at least three biological replicates.

FabT represses the promoter P*fabK*, which is believed to drive the expression of the genes from the second FAS-II operon (*fabK*-*acc*A). However, to our knowledge the presence of other promoters, or specific processing sites in this operon, have not been described. In our conditions, a full-length transcript corresponding to *fabK*-*acc*A was not detected, and all the tested *fab* transcripts from this polycistronic message could be found alone. In rare instances, we could detect a message with about 6 kb (data not shown). This might indicate the existence of some additional processing events, controlling the expression of these genes post-transcriptionally. In confirmation of the RNA-Seq, these results show an accumulation of all the *fab* transcripts evaluated here in the absence of RNase R (apart from *fabM*). In all cases the WT levels were partially restored in the complemented strain (Δ*rnr*+R), suggesting a role for this enzyme in the turnover of these mRNAs.

To further demonstrate the direct role of RNase R in this process, we have measured the stability of these messages in the WT and in the RNase R mutant. The results of the decay assays are shown in Figure 2B and the respective plots are presented in Figure 2C. The presence of RNase R is shown to significantly decrease the stability of all the genes tested (Table I). The half-life of the *fabT*-*fabH* operon decreased by about 6 min when the probe specific for *fabH* was used, and only ≈3 min with the *fabT*-specific probe. Degradation of *fabK, fabG* and *fabF* messages is also faster in the presence of RNase R leading to differences in the stability of about 3.5, 1.7 and 3.2 min, respectively between these strains. Surprisingly, the transcriptional unit *fabT-fabH* presents different stabilities depending on the probe used in the Northern experiments. However, RNase R degrades RNA in the 3’ to 5’ direction and *fabH* is located downstream of *fabT* nearer the 3’ end of the transcript. Thus, it is not surprising that *fabH* is the first message to be degraded, which results in a higher difference between the WT and *Δrnr*. Alternatively, it might be possible that degradation of the transcriptional unit also involves internal cleavage thus generating differences in half-life between fabT and fabH.

**Table I.** Half life of *fab* transcripts in the WT and Δ*rnr*.

| Strain \ Gene | *fabT* | *fabH* | *fabK* | *fabG* | *fabF* |
| --- | --- | --- | --- | --- | --- |
| WT | 1.13 min | 0.92 min | 1.49 min | 1.32 min | 1.26 min |
| $\Delta rnr$ | 3.29 min | 5.92 min | 5.25 min | 2.26 min | 4.08 min |

Overall, this data is in agreement with a direct role of RNase R in the modulation of the expression level of the FAS-II genes cluster, by controlling the turnover of the respective transcripts.

### Impact of RNase R on the Membrane Fatty Acid Composition

Alterations in the expression level of genes from the *fab* cluster have long been known to affect the fatty acid balance leading to abnormal composition of cell membranes [14, 19]. Thereby we decided to determine the fatty acid composition of the WT and Δ*rnr* membranes. To this end, the fatty acid composition was determined by gas chromatography. As shown in Figure 3A we could observe the presence of the following saturated fatty acids: lauric acid (C12:0), myristic acid (C14:0), palmitic acid (C16:0) and stearic acid (C18:0) in both strains. Regarding unsaturated fatty acids we identified mono-unsaturated FA, including two C16:1 isomers (Δ7 and Δ9), three C18:1 isomers (Δ9, Δ11 and Δ13) and the di-unsaturated FA C18:2, in agreement with the literature [20, 21]. Although the total fatty acid content in the mutant cells was similar to the levels measured in the WT (Figure 3B), the Δ*rnr* strain presented alterations in their relative proportions. Mutant cells had lower percentages of C14:0 and C16:0 and increased proportions of all species of C18 FA (C18:0, the three C18:1 isomers, as well as 18:2) (Figure 3A). These changes in individual fatty acids led to alterations in derived parameters related to fatty acid elongation and unsaturation processes, such as the ratio C18/C16 (Figure 3C), which was higher in the mutant, and the proportion of saturated fatty acids (∑Sat) (Figure 3D) which was lower in the same strain. These results suggest that the absence of RNase R affects FA biosynthesis and unsaturation, leading to an altered membrane fatty acid composition.

**Figure 3.**
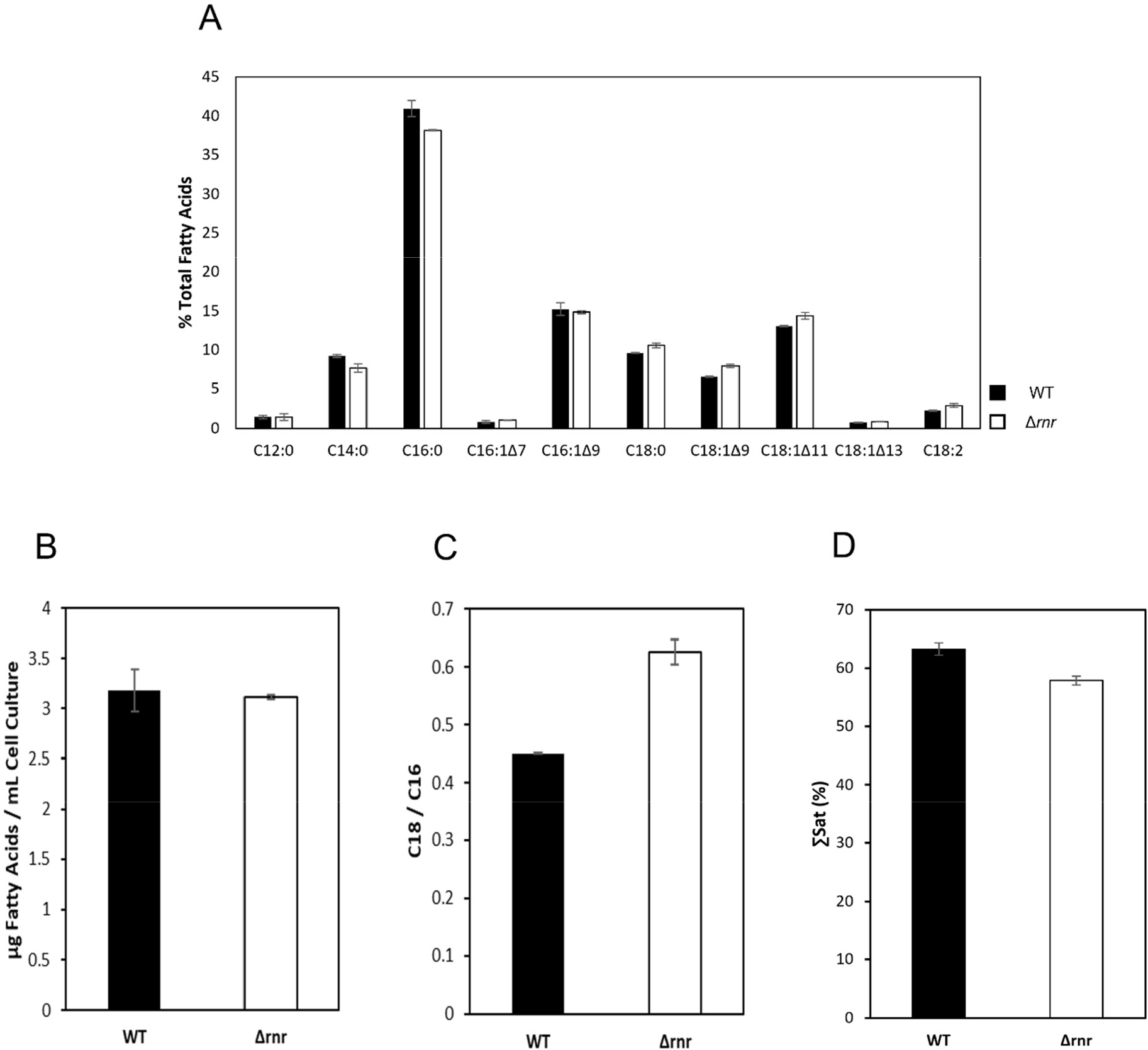
Influence of RNase R on the fatty acids of *S. pneumoniae*. **A)** Fatty acid composition of wild type (WT) and mutant (*Δrnr*) cells of *S. pneumoniae*, expressed as percentage of total fatty acids. **B)** Total fatty acids contents (expressed in μg FA/mL cell culture). **C)** Ratio between FA with 18 carbons to FA with 16 carbons. **D)** Percentage of saturated fatty acids (∑sat). Cultures of *S. pneumoniae* strains were grown to an OD_600_ ≈ 0.3. Fatty acid methyl esters prepared from bacterial cells were analyzed by gas chromatography. Values correspond to average ± standard deviation, n=3. These experiments were performed in triplicate.

### Effects on resistance to stress conditions

Composition of the membrane lipid bilayer determines its biophysical properties and affects membrane integrity and permeability [22]. Membrane FA composition is influenced by different stress conditions and has been related to stress tolerance, namely temperature, pH, oxidative stress and detergent lysis [13, 19, 21, 23, 24]. We have examined the sensitivity of the wild type, Δ*rnr* and Δ*rnr*+R to detergent lysis and have observed an increased sensitivity of the Δ*rnr* mutant to sodium deoxycholate when compared to the strains containing RNase R (Figure 4A). Changes in membrane fluidity caused by the lack of RNase R might be the reason for the decreased tolerance to sodium deoxycholate observed [13, 19] in the mutant strain. Bacterial adaption in response to oxygen and oxidative stress includes changing the fatty acid composition of the membrane and this is also documented in *S. pneumoniae* [14, 19, 21, 23]. The data presented here shows that membrane lipids are affected by RNase R and we have already demonstrated that this enzyme is essential to overcome the growth defects caused by H_2_O_2_ [4]. This may well be related with a role for this ribonuclease on the regulation of membrane homeostasis under oxidative stress. In this case, we might expect to find an increased expression level of RNase R in cells challenged with H_2_O_2_. To test this hypothesis, we compared the RNase R protein levels by Western blot before and after 5 mM H_2_O_2_ addition (Figure 4B). At 30 min post-exposure we observed about 50% increase in the RNase R levels. Elevation of RNase R levels after oxidative stress challenge provides additional evidence for an important role of this enzyme in the adaptive response of the pneumococcus to oxidative stress. Taken together, this may be ascribed to defects in membrane adaptation due to the RNase R role on the modulation of the FAS-II cluster expression.

**Figure 4.**
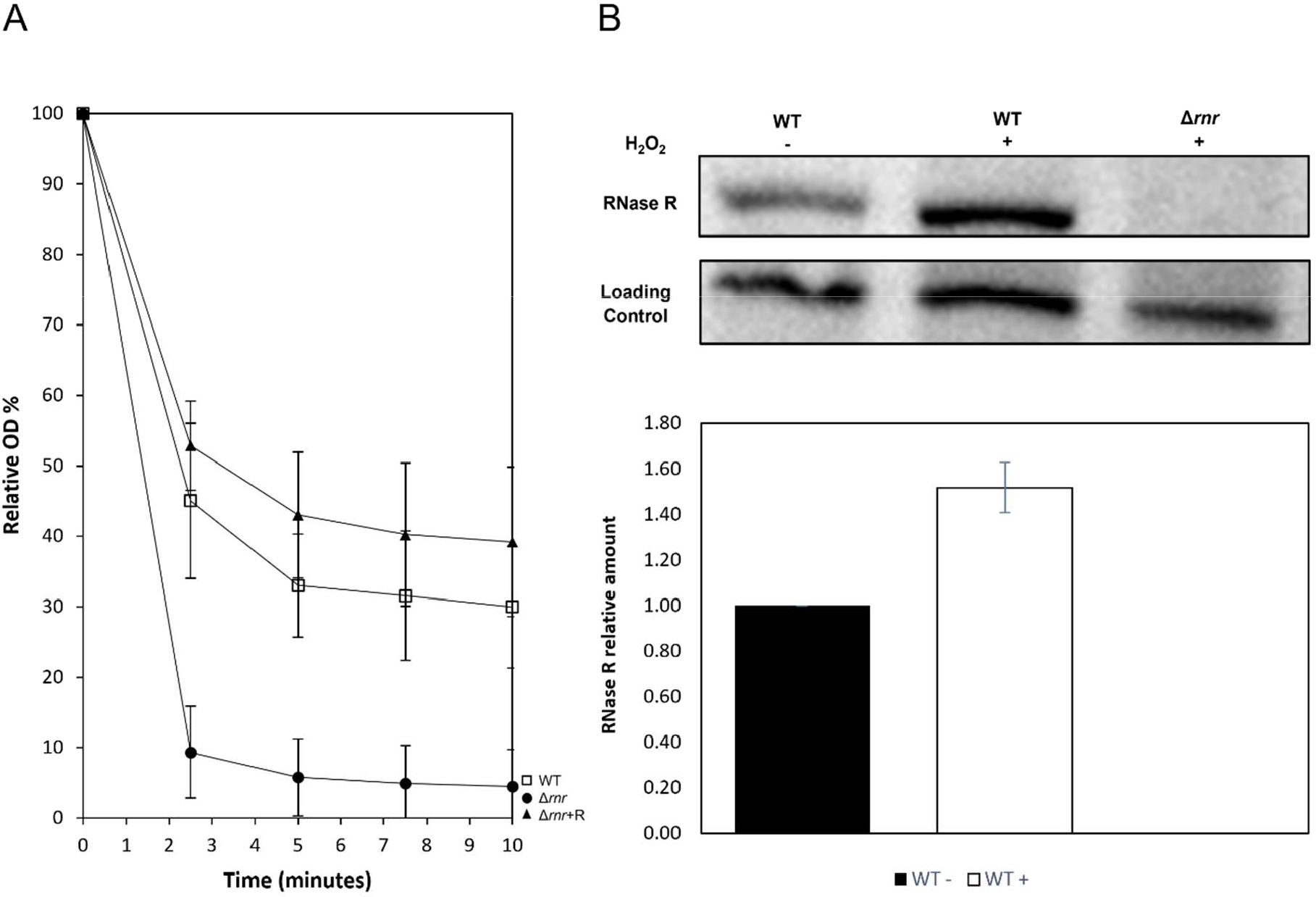
RNase R implications on stress tolerance. A) Tolerance to detergent lysis. Relative cell density in the presence of 2.5% sodium deoxycholate. WT, Δ*rnr*, and Δ*rnr*+R strains were grown in THY medium, until the beginning of the exponential phase (OD_600_≈0.3-0.4) and then challenged with 2.5% sodium deoxycholate. The absorbance at 600 nm was monitored for the indicated time-points. The results were normalized to the optical density obtained for the same culture without detergent. These results were obtained from at least three independent essays. **B) RNase R levels increase in the presence of H**_**2**_**O**_**2**_. Expression of RNase R was analysed in the wild type (WT) by Western blot before (-) and after 5 mM H_2_O_2_ addition (+) as indicated on top of the image. Δ*rnr* mutant was used as negative control. A non-specific band (Control) detected with the same antibodies was used as loading control. 100 μg of total protein were separated in a 7% tricine-SDS-polyacrylamide gel (Haider *et al*., 2010) and blotted into a nitrocellulose membrane. Immunodetection was performed with RNase R-specific antibodies using Western Lightning Plus-ECL Reagents (PerkinElmer) and the iBright1500 (Invitrogen by Thermo Fisher Scientific) was used to obtain the images. **C) RNase R affects colony morphology**. Bacterial cells were grown in THY medium until OD_600_ ≈ 0.3. Equal aliquots of the different strains were plated in THY agar medium supplemented with 5 % blood and plates were incubated 24h at 37°C in a 5 % CO_2_ atmosphere. Bacterial growth was compared in the three strains (WT, Δ*rnr* and Δ*rnr*+R). Images were obtained with Carl Zeiss Axis Zoom microscope (16x magnification).

## Discussion

Maintaining cytoplasmic membrane integrity is critical for bacterial survival and adaptation to stress conditions, and bacterial response to different environmental conditions involves adjusting its lipid composition [27]. Changes in membrane lipid composition result in alterations in its biophysical properties, and a correct FA profile is essential for membrane homeostasis [20] reviewed by [15]. In this work we show that RNase R plays an important role in the control of *S. pneumoniae* membrane composition, by regulating the amount of several *fab* genes, possibly influencing the stress response ability of this pathogen.

Global transcriptomic analysis of an RNase R mutant strain highlighted that RNase R affects several different transcripts with different roles in the cell. We found that the majority of the transcripts affected by RNase R deletion were from ABC transporters and PTS systems. A previous study on the transcriptomic differences between aerobic and anaerobic growth of *S. pneumoniae* had also found several transcripts from ABC transporters and PTS systems to be affected [28]. Furthermore, the authors found that when comparing the aerobic and anaerobic transcriptomes oxidative stress genes were also affected. Our study was conducted under anaerobic conditions and the similarities between the results obtained in our study and the Bortoni et al work [28] could suggest that RNase R might have a role in the anaerobic metabolism. In fact, there are also some anaerobic related transcripts that we found to be upregulated in the RNase R mutant, namely *nrdG* and *nrdD* (Table S1).

It specifically came to our attention the elevated expression of most of the genes belonging to the FA biosynthesis pathway detected in the absence of RNase R. *S. pneumoniae* adopts the FAS-II synthesis pathway, in which membrane fatty acids are produced by a set of enzymes encoded by the genes of the FAS-II cluster. The initiation enzyme FabH, performs the first condensing reaction on the malonyl-ACP supplied by the concerted action of the multisubunit acetyl-Coenzyme A carboxylase (AccABCD) and the manolyl-ACP transacylase (FabD). The nascent fatty acids are extended by the elongation module two carbons at a time, in a series of repetitive steps performed in *S. pneumoniae* by the enzymes FabG, FabZ, FabK and FabF (reviewed in [15, 29]). The end products of the pathway consist primarily of 16- and 18-carbon acyl-ACPs, which can be saturated (SFA) or unsaturated (UFA) and consist of the hydrophobic portion of the membrane phospholipids. FabM introduces the double bond and the UFA/SFA ratio ultimately depends on the competition between FabM and FabK for the substrate generated by FabZ [15, 30]. The increasing level of most of *fab* genes revealed by RNA-Seq, was confirmed by the steady-state analysis of *fab* mRNA transcripts. Further decay studies showed a global stabilization of these transcripts in the absence of RNase R. This is supportive of a role for RNase R in the degradation of this polycistronic message, thereby regulating the final amount of these transcripts in the cell. Alterations in the expression of these genes are known to result in a different fatty acid composition of membrane phospholipids. Overexpression of all genes associated with chain elongation resulted in longer FA chains [13, 14]. We observed increased expression levels of genes from both the initiation and elongation modules, which is in agreement with the increased long-chained FA observed in the Δ*rnr* membrane. Notably, FabG and FabF, whose transcripts are stabilized in the strain lacking RNase R, might play a role in the chain length of fatty acids. Overexpression of the essential condensing enzyme FabF was shown to increase the abundance of C18 species as compared to C16 [19]. The FabG reductase is involved in the elongation of FA chains by catalysing the reduction of the carbonyl group from the substrates produced by the condensing enzymes (whether FabH or FabF) (reviewed by [15]). Since the length of the fatty acid chain is ultimately determined by the number of cycles of condensation and reduction that occur during the elongation process, increased FabG levels might presumably lead to long-chained FA.

It could be expectable that the global increased expression of *fab* genes would result in higher amounts of FA. However, our results indicate similar amounts of total FA in the WT and Δ*rnr* membranes. These results are consistent with the fact that we did not observe variations in the *acpP* level, which encodes the acyl carrier protein (ACP). This protein is a central player in the FAS-II pathway that carries all of the FAS-II-pathway intermediates and is fundamental for the fatty acid unit starter production.

Transcriptional regulation of the FAS-II cluster is under the control of FabT, a transcriptional repressor that binds both promoters of the cluster, regulating its own transcription and the expression of all of *fab* genes, except *fabM* [13, 16]. A putative FabT binding site was also reported upstream of *fabM* [30] and recent works have shown a FabT-dependent regulation of *fabM* [27, 31]. Since FabT affinity for DNA is increased by binding to long chain acyl-ACPs [16], *fabM* repression was proposed to rely on the formation of highly repressive FabT complexes formed with longer acyl-ACPs, which were present in the conditions tested by the authors [31]. In our case however, the increased levels of *fabT* resulting from RNase R elimination do not seem to affect the expression level of *fabM*. As a repressor, overexpression of FabT was expected to reduce the expression of all the *fab* genes, with the possible exception of *fabM*. On the contrary, our observations showed that *fabT* levels increased concomitantly with the other *fab* genes in the Δ*rnr* mutant. The fact that FabT auto-regulates its own expression might explain the higher expression levels of the other genes of the cluster in the Δ*rnr* mutant. Indeed, this result is in accordance with a previous observation that *fabT* overexpression did not repress *fabK* below the WT levels [13, 19].

Differences in the ratio of unsaturated to saturated FA are determined in *S. pneumoniae* by the direct competition between FabM, and independent overexpression of FabK was previously shown to raise the proportion of SFA [13]. Increased levels of FabK were also detected in the absence of RNase R, but in our study, we observed higher levels of UFA. However, the increased levels of *fabK* in Δ*rnr* were accompanied by the increment of other *fab* genes, probably altering the amount of available substrate. In fact, Mohedano et al [14] did not detect variation in the ratio of unsaturated to saturated FA when several enzymes of the pathway were simultaneously increased.

The data presented here is suggestive of an important role of RNase R in the pneumococcal membrane homeostasis. In the absence of this enzyme, the proportion of membrane FA changed from short-chained saturated FA to long-chained unsaturated FA, which is suggestive of a less rigid membrane in the Δ*rnr* mutant cells. The higher fluidity of the mutant membrane might be responsible for its higher sensitivity to detergent. Alterations in the membrane homeostasis caused by deletion of *fabT*, were also shown to significantly increase sensitivity to deoxycholate [13]. This strain also showed an acid-sensitive growth phenotype and pH changes are known to induce alterations in the FA content of the membrane [13, 21, 31-33]. Loss of RNase R, however, did not increase sensitivity to acidic pH (data not shown).

Overall, maintaining membrane homeostasis is critical for cell survival under stress conditions, and the ability to adapt to these conditions is determined in large part by the fatty acid composition and the capacity of bacteria to perform the necessary adjustments (reviewed by [29]). In *S. pneumoniae*, a significant alteration in the membrane FA profile was related to increasing amounts of endogenous H_2_O_2_, which was shown to inhibit FabF activity by oxidizing its active site [21, 23]. Higher levels of active FabF, observed under anaerobic growth conditions, correlated with a considerable enhancement in FA unsaturation together with an increase in C18:C16 ratio. Interestingly these results mostly resemble the membrane composition changes observed after elimination of RNase R, which also led to a phenotype characterized by high proportions of long, unsaturated fatty acyl residues [21, 23]. Changes in the pool of membrane lipids in response to transitions between aerobic and anaerobic growth have long been reported to occur in *Staphylococcus aureus* [34]. The essential regulator YycFG, shown to regulate membrane composition by inhibiting FabT transcription, might also respond to oxygen, as previously suggested [17]. We have previously shown that deletion of RNase R strongly affected the capacity of the deleted strain to cope with oxidative stress [4]. Here we show that RNase R expression is induced by H_2_O_2_, evidencing an important role of this ribonuclease under these conditions. Alterations of membrane lipid fatty acyl chains induced by the lack of RNase R might be, at least in part, responsible for the increased mortality observed in *Δrnr* in the presence of H_2_O_2_ [4]. Interestingly our RNA-Seq results also pinpoint two genes related to oxidative stress, which were upregulated in Δ*rnr*. In this strain, the absence of RNase R might compromise the expression levels of the FAS-II genes required to perform the necessary adjustments in the membrane lipids pool during adaptation to oxidative stress.

The cell membrane is an essential component of the bacterial cell, acting as a protective barrier from the external environment. It is thus crucial for adaptation to environmental changes. The existence of several regulatory mechanisms that control the expression of the FAS-II genes and regulate the activity of some enzymes of the pathway is essential in allowing the cells to maintain a proper membrane fluidity. This work extends the list of known regulators by identifying RNase R as one intervenient in the control of the expression levels of the FAS-II cluster, which in turn determines the FA content of the membrane.

## Materials and Methods

### Bacterial Strains, Oligonucleotides and Growth Conditions

*S. pneumoniae* strains were grown in Todd Hewitt medium, supplemented with 0.5% yeast extract (THY) at 37°C without aeration, or in THY agar medium supplemented with 5% sheep blood (Thermo Scientific) at 37°C in a 5% CO_2_ atmosphere. When required growth medium was supplemented with 3 μg/ml chloramphenicol (Cm), 1 μg/ml erythromycin (Ery) or 150 μg/ml kanamycin (Km) as specified.

All *S. pneumoniae* strains are isogenic derivatives of the JNR7/87 capsulated strain – TIGR4 and are listed in Table S1 (Supplementary Material).

DNA sequencing was performed at the genomic central service from Centro Nacional de Microbiología (CNM, ISCIII) and oligonucleotide synthesis was performed at CNM, ISCIII and STAB Vida. Oligonucleotide sequences are listed in Table S2 (Supplementary Material).

### Construction of *Δrnr*+R strain of *S. pneumoniae*

To construct the *Δrnr+R* strain, an ectopic copy of *rnr* gene was introduced into the Δ*rnr* strain chromosome within the *bgaA* locus through allelic replacement [35]. For this purpose, four fragments were generated by PCR (named from 1 to 4). PCR 1 and 4 contained 5’- (coordinates 615828..616648) or the 3’-sequence (620648..621431) of *bgaA* locus and were amplified from the TIGR4 genome using primer pairs BgaA_F/BgaA_Bam and BgaA_Nhe/BgaA_R, respectively. PCR 2 corresponded to the Km resistance cassette and was amplified from pR410 plasmid [36] using primers Km_Bam and Km_Kpn. The PCR 3, containing the *rnr* gene fused to its natural Psec promoter, was amplified from the TIGR4 genome using primers Psec_rnr (harbouring the promoter sequence) and RNR_Nhe. The four PCR fragments were purified, digested with the corresponding restriction enzyme (target sites included in the oligonucleotides), mixed together, and ligated with T4 DNA ligase. A final PCR reaction (PCR 5) was performed using primers BgaA_F2 and BgaA_R2 and the ligation mix as DNA template. The resulting 5075-Kb fragment containing the up sequence of *bgaA*, the Km resistance cassette, the Psec-*rnr*, and the down *bgaA* sequence (Figure S3) was used to transform the *Δrnr* as previously described [37]. Transformants were selected with 250 µg/mL km and positive clones were confirmed by sequencing.

### RNA Extraction from *S. pneumoniae* Cultures

Overnight cultures of *S. pneumoniae* TIGR4 WT and derivatives were diluted in pre-warmed THY to a final OD_600_ of 0.05 and incubated at 37°C until OD_600_ ≈ 0.3. 25 ml of culture was collected, mixed with 1 volume of stop solution (10 mM Tris pH 7.2, 25 mM NaNO_3_, 5 mM MgCl2, 500 µg/ml Cm) and harvested by centrifugation (10 min, 6000 x *g*, 4°C). For decay studies, transcription was stopped by addition of rifampicin (0.5 mg/ml) and nalidixic acid (20 µg/ml) to growing cells (OD_600_ ≈ 0.3). At the indicated times, 25 ml of culture was collected, mixed with 1 volume of stop solution and harvested by centrifugation in the conditions described above. In both cases, total RNA was extracted using Trizol reagent (Ambion) essentially as described by the manufacturer with some modifications as previously described [11]. RNA concentration was estimated using a Nanodrop 1000 machine (Nanodrop Technologies). RNAs used for RNA-Seq were further checked using Bioanalyzer (Agilent) for RNA integrity determination.

### Northern blot Analysis

For Northern blot analysis, total RNA samples were separated under denaturing conditions by 1.5 % agarose MOPS/formaldehyde gel. RNA was transferred to Hybond-N+ membranes (GE Healthcare) by capillarity using 20× SSC as transfer buffer, and UV cross-linked to the membranes immediately after transfer. RNAs integrity and loading control were accessed by staining the membranes in a methylene blue solution (0.03 % methylene blue 0.3 M CH_3_CO_2_Na pH 5.2) and distaining in water. Membranes were then thoroughly washed in 0.5%, 1% SDS and water, and then hybridized in PerfectHyb Buffer (Sigma) for 16 h at 68°C for riboprobes and 43°C in the case of oligoprobes. After hybridization, membranes were washed as described [38]. Signals were visualized by Phosphorimager (TLA-5100 Series, Fuji) and analysed using the ImageQuant software (Molecular Dynamics).

### Hybridization Probes

PCR fragments used as templates for riboprobes were amplified with DreamTaq (Thermo Fisher) and purified using the “PCR Clean-up System” (Macherey-Nagel).

Riboprobe synthesis and oligoprobe labelling were performed as previously described [38]. PCR reactions were carried out using the following primer pairs: smd251/smd252, smd253/smd254, smd075/smd100, smd248/smd249, smd079/smd102 and smd188/smd189 respectively for *fabT, fabH, fabK, fabG, fabF and fabM* specific probes. DNA probes for 23S rRNA were generated using oligonucleotides cbr014, labeled at its 5’-end with [^32^P]-γ-ATP using T4 Polynucleotide kinase (Thermo Fisher) according to the supplier instructions.

### RNA Seq

Total RNA was sent to StabVida (Portugal) for sequencing (Illumina, paired-end, 150bp, 20M reads). rRNA was depleted from the samples with the RiboZero kit and sequencing libraries were constructed with the TruSeq kit (Illumina) for bacteria. The quality of the RNA-Seq data was confirmed using the fastQC program. Contaminant adapters and low-quality reads were removed with cutadapt [39] and the remaining reads were mapped against *S. pneumoniae* TIGR4 genome (NC_003028.3, downloaded from NCBI genome database) using Bowtie2 program [40]. Samtools [41] was used to sort the mapping files by genomic position and quantification of RNA-Seq data was performed using FeatureCounts [42]. Transcripts expression was calculated using the RPKM normalization method. Differences in expression between the two strains was obtained by calculating the Log_2_Fold change (Log_2_FC). Results were filtered to have differences above 0.5 and below -0.5 of Log_2_FC and to have RPKM values above 5 in at least one of the samples. Furthermore, all hypothetic proteins were removed from the final analysis.

### Fatty acid composition analysis

Overnight cultures of S. pneumoniae strains were diluted in prewarmed THY to a final OD_600_ of 0.05 and incubated at 37°C until OD_600_ ≈ 0.3. 40 ml of culture were collected, mixed with 1 volume of stop solution (described above) and harvested by centrifugation (10 min, 6000 x *g*, 4°C). Bacterial pellets were stored at -80°C until further analysis.

Fatty acid profiling was performed by direct *trans*-esterification of bacterial pellets in methanol-sulfuric acid (97.5%, v/v) at 70◦C for 60 min, as described in [43], using margaric acid (C17:0) as internal standard. Fatty acid methyl esters (FAMEs) were recovered with petroleum ether, dried under an N_2_ flow and re-suspended in hexane. One microliter of the FAME solution was analysed in a gas chromatograph (Varian 430-GC gas chromatograph) equipped with a hydrogen flame ionization detector set at 300◦C. The temperature of the injector was set to 270◦C, with a split ratio of 50. The fused-silica capillary column (50 m × 0.25 mm; WCOT Fused Silica, CP-Sil 88 for FAME; Varian) was maintained at a constant nitrogen flow of 2.0 mL min-1, and the oven temperature was set at 190◦C. FA identification was performed by comparing retention times with standards (Sigma-Aldrich), and chromatograms were analysed by the peak surface method using the Galaxy software.

### Lysis Assay with Sodium Deoxycholate

*S. pneumonia* cultures were grown in THY medium to a density of mid-log phase and then diluted to OD_600_ ≈ 0.05. At OD_600_ ∼ 0.3 bacteria were challenged with 2.5% of sodium deoxycholate. OD_600_ of the treated cultures was then followed 2.5, 5, 7.5 and 10 minutes after sodium deoxycholate addition. As negative control the optical density of the same bacterial cultures without sodium deoxycholate was also monitored.

### Total protein extraction and Western immunoblotting

Early exponential phase cultures of *S. pneumoniae* TIGR4 WT (OD at 600 nM ≈ 0.3–0.4) were challenged with 5 mM of hydrogen peroxide and incubated at 37 ◦C. 20 ml culture samples were collected before and 30 min after stress challenge, mixed with 1 volume of stop solution (defined above) and harvested by centrifugation (10 min, 6000 x *g*, 4°C). The cell pellet was resuspended in 100 µl of TE buffer supplemented with 1 mM PMSF, 0.15% sodium deoxycholate and 0.01% SDS. After 15 min incubation at 37°C, SDS was added to a final concentration of 1%. Protein concentration was determined using a Nanodrop 1000 machine (NanoDrop Technologies). Total protein samples were separated in a modified 7% tricine-SDS-polyacrylamide gel as described [44].

After electrophoresis, proteins were transferred into a nitrocellulose membrane (Hybond ECL, GE Healthcare) by electroblotting using the Trans-Blot SD semidry electrophoretic system (Bio-Rad). The membrane was probed with 1:500 dilution of anti-RNase R antibodies [10]. ECL anti-rabbit IgG peroxidase conjugated (Sigma) was used as the secondary antibody in a 1:10000 dilution. Immunodetection was conducted via a chemiluminescence reaction using Western Lightning Plus-ECL Reagents (PerkinElmer).

## Acknowledgments

We are thankful to Teresa Baptista da Silva for technical assistance.

