## Supplementary Material for "RNase R Controls Membrane Fatty Acid Composition in *Streptococcus pneumoniae*"

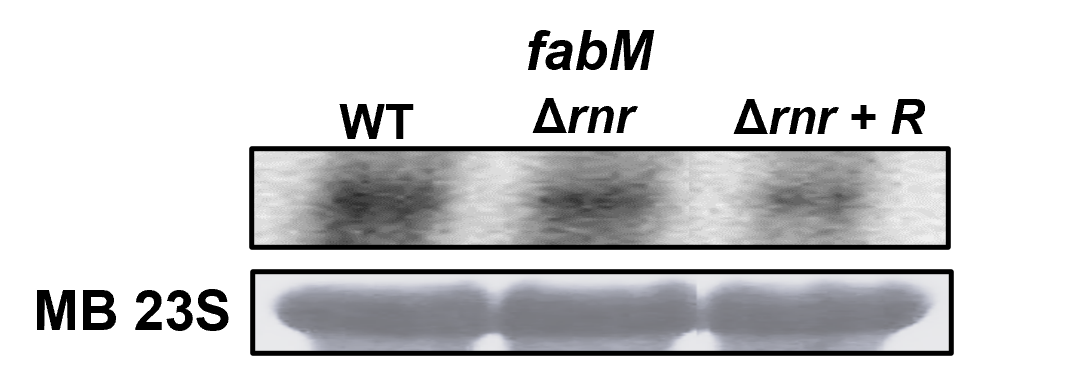

**Figure S1 – Expression of *fabM* is not affected by RNase R.** Northern blot analysis of total RNA samples extracted from the wild type (WT), RNase R mutant (Δ*rnr*) and Δ*rnr* expressing RNase R *in trans* (Δ*rnr*+R). 20 µg of total RNA were separated on 1.5 % agarose gels, transferred to a Hybond-N+ membrane and hybridised with a specific probe for *fabM* mRNA. Loading control was performed by staining the membrane with methylene blue (MB) before hybridization with the probe.

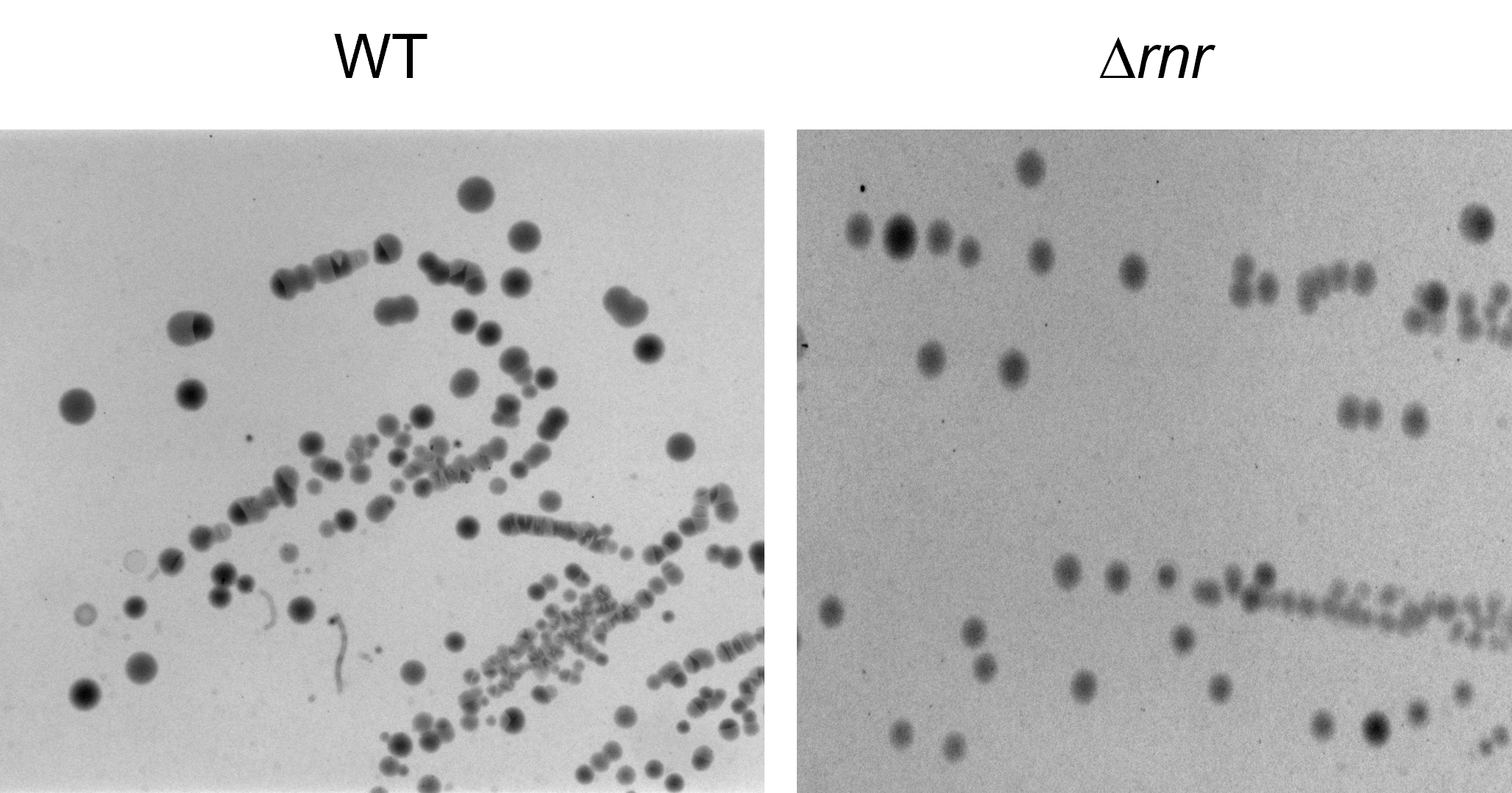

**Figure S2 – Phase variation is not affected by RNase R.** WT and Δ*rnr* exponentially grown cultures were plated on blood agar with appropriate antibiotics, supplemented with 6300 U of Catalase. Plates were incubated at 37°C in a 5 % CO_2_ atmosphere and after ≈ 24h of incubation bacterial colonies were observed using Carl Zeiss Axis Zoom microscope.

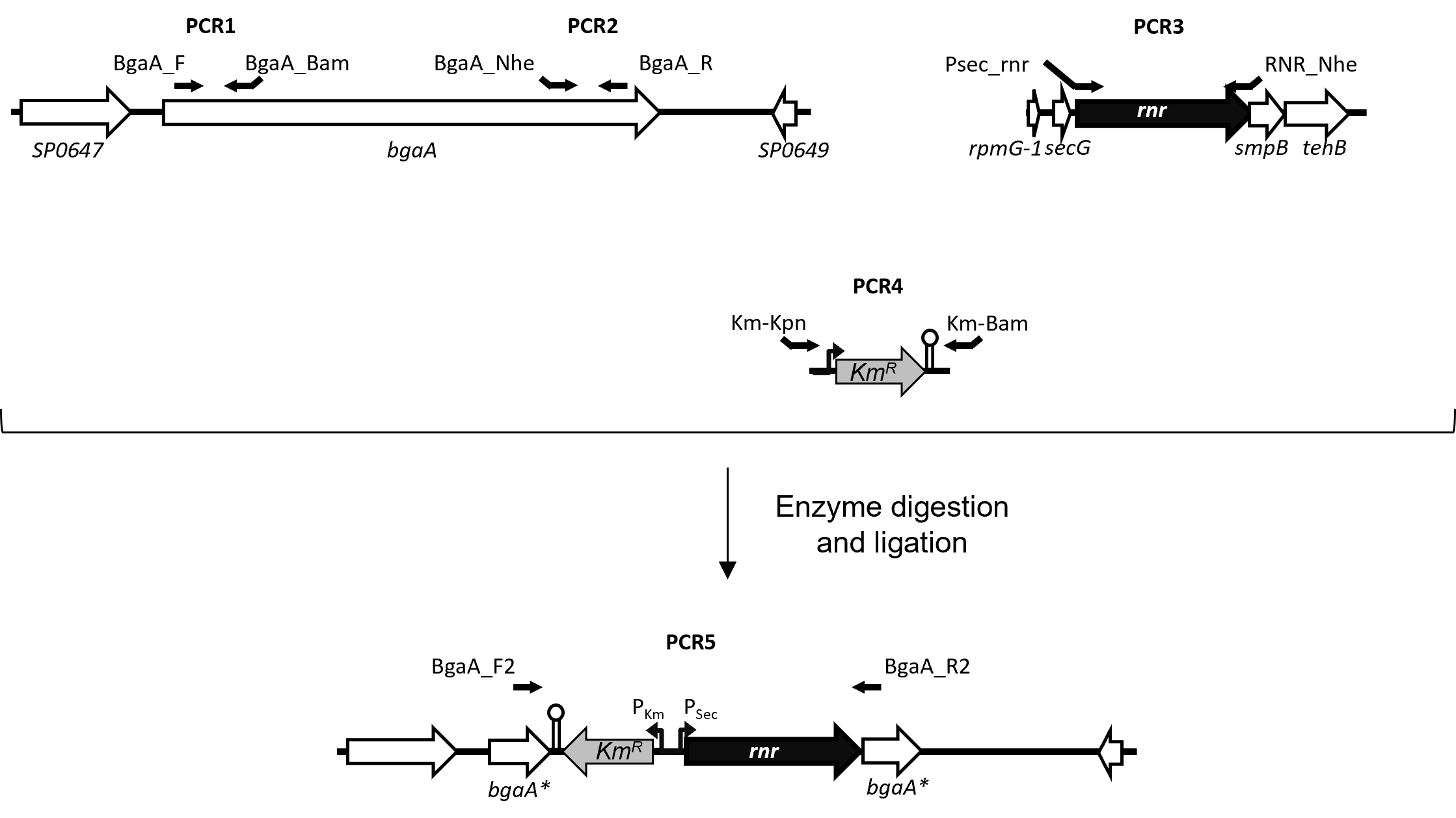

**Figure S3 –** Schematic representation of the Δ*rnr*+R strain construction**.**

**Table S1 –** List of strains used in this work.

| Strain | Relevant characteristics | Reference |
| --- | --- | --- |
| *S. pneumoniae* |  |  |
| JNR7/87 (TIGR4) |  | [1] |
| Δ*rnr* | TIGR4 *rnr*^−^ (Cm^R^) | [2, 3] |
| Δ*rnr*+R | Δ*rnr* *bgaA*::*rnr-kan^R^* expressing RNase R (Cm^R^, Kan^R^) | [This work] |

Kan^R^: Kanamycin resistant; Cm^R^: Cloramphenicol resistant

| Oligo name | Nucleotide sequence (5’ to 3’) |
| --- | --- |
| smd251 (*fabT*) | gttttttttttaatacgactcactataggGAAGCGTTTATGCAGTCTATG |
| smd252 (*fabT*) | CATCGGTAAGGCTCCAGACG |
| smd253 (*fabH*) | gttttttttttaatacgactcactataggCTTGGCTACATCTCGAATGGC |
| smd254 (*fabH*) | CTTAGCTGAGAGTCTCAATAG |
| smd100 (*fabK*) | gttttttttttaatacgactcactataggACAAGCCCTGCGATTTGACC |
| smd75 (*fabK*) | GTGGAAGACATCGTGGATCTC |
| smd248 (*fabG*) | GTCAACAATGCAGGGATTAC |
| smd249 (*fabG*) | gttttttttttaatacgactcactataggATGATAGCACCTTCTCTGGC |
| smd079 (*fabF*) | GTAGCCATGCGTTTTGGTGC |
| smd102 (*fabF*) | gttttttttttaatacgactcactataggGACGCATAGCTTCGATGGTG |
| smd188 (*fabM*) | gttttttttttaatacgactcactataggTCTGCCGTTAAAGCCTCTCC |
| smd189 (*fabM*) | CGCAGCGAATATGGCTGTTG |
| cbr014 (23S rRNA) | GCTCTACCTCCAAGAATCTC |
| BgaAF | ACCGCTCAAGGAAGATGCTA |
| BgaA_Bam | CGC**GGATCC**CGTACAAGGCAGGTTTGTC |
| BgaA_Nhe | GCCGC**GCTAGC**CAGTTGCTGCGGTTAAAC |
| BgaA_R | CTTGGTGCAAGGAAGGTCAT |
| Km_Bam | GCGC**GGATCC**ATCGATACAAATTCCTC |
| Km_Kpn | GCGC**GGTACC**AAGGGCCCGTTTGATTTTTAATG |
| Psec_rnr | GCGC**GGTACC**TACATCTTGAATCTAAGTAGATTTTATGGTAAAATAATAGGAATCTAGAAAGAAAATATGAAAGATAGAATAAAAG |
| RNR_Nhe | CGGCG**GCTAGC**TTTCTGCAGTTTATTTTGTGCG |
| BgaA_F2 | GAAGGTGGACAGCTCAACGGTG |
| BgaA_R2 | CTCAATCATCGCAACACCGCTCTTATC |

**Table S2 –** Oligonucleotides used in this work.
